# Maize pan-transcriptome provides novel insights into genome complexity and quantitative trait variation

**DOI:** 10.1101/022384

**Authors:** Minliang Jin, Haijun Liu, Cheng He, Junjie Fu, Yingjie Xiao, Yuebin Wang, Weibo Xie, Guoying Wang, Jianbing Yan

**Author notes:** Correspondence should be addressed to J.Y. and J.F. These authors contributed equally to this work (Minliang Jin, Haijun Liu, Cheng He).

## Abstract

Variation in gene expression contributes to the diversity of phenotype. The construction of the pan-transcriptome is especially necessary for species with complex genomes, such as maize. However, knowledge of the regulation mechanisms and functional consequences of the pan-transcriptome is limited. In this study, we identified 13,382 nuclear expression presence and absence variation candidates (ePAVs, expressed in 5%~95% lines; based on the reference genome) by re-analyzing the RNA sequencing data from the kernels (15 days after pollination) of 368 maize diverse inbreds. It was estimated that only ~1% of the ePAVs are explained by DNA sequence presence and absence variations (PAV). The ePAV genes tend to be regulated by distant eQTLs when compared with non-ePAV genes (called here core expression genes, expressed in more than 95% lines). When the expression presence/absence status was used as the “ genotype” to perform genome-wide association study, 56 (0.42%) ePAVs were significantly associated with 15 agronomic traits and 1,967 (14.74%) with 526 metabolic traits, measured from the mature kernels. While the above was majorly based on the reference genome, by using a modified ‘assemble-then-align’ strategy, 2,355 high confidence novel sequences with a total length of 1.9Mb were found absent in the current B73 reference genome (v2). Ten randomly selected novel sequences were validated with genomic PCR. A simulation analysis suggested that the pan-transcriptome of the maize whole kernel is approaching a maximum value of 63,000 genes. Two novel validated sequences annotated as NBS_LRR like genes were found to associate with flavonoid content and their homologs in rice were also found to affect flavonoids and disease-resistance. Novel sequences absent in the present reference genome might be functionally important and deserve more attentions. This study provides novel perspectives and resources to discover maize quantitative trait variations and help us to better understand the kernel regulation networks, thus enhancing maize breeding.

## INTRODUCTION

Maize shows an amazing degree of phenotypic variation due to the outcrossing nature, and to natural and artificial selection during the rapid worldwide population expansion (Yan et al. 2011). Phenotypic variation has been explored by QTL mapping and genome-wide association studies (GWAS; Huang and Han 2014). As it becomes clear that the differences in transcript abundance are a major contributor to phenotypic evolution (Cubillos et al. 2012; Albert and Kruglyak 2015; Liu et al. 2015), allelic variation effects on the transcriptome, which reflect both genetic and epigenetic regulation, should be explored at a genome-wide level (Fu et al. 2013).

Presence/absence genomic sequence variation (PAV) is important in reshaping individual performance (Springer et al. 2009). PAV at the genomic level would be reflected in the transcriptome ePAV (expression Presence and Absence Variation). The ePAV not only reflect genomic structural variation, but also the variations in genetic and epigenetic regulatory elements. Thus it is essential to characterize the ePAV genes and their possible functions.

Most genome-wide genetic studies focus the genetic elements present in the reference genome. It is now recognized that a portion of the genomic content is only present in a subset of individuals within a species, (termed the dispensable genome; Medini et al. 2005) especially in diverse species, such as maize. The genome-wide comparison between B73 and Mo17 (Springer et al. 2009) and within an expanded panel including teosinte (ancestral maize) lines (Swanson-Wagner et al. 2010) demonstrated that a considerable portion of the genome (~50%) was not shared. The widespread dispensable genes, i.e those showing present/absent variation, have been proposed to be important for phenotypic diversity in inbred collections and for heterotic performance in hybrids (Lai et al. 2010; Hansey et al. 2012).

The rapid development of next generation sequencing technology and the decrease in cost provide us an opportunity to sequence many individuals within a species to build up the pan genome, or the sequences which, taken as a whole from all individuals, define a species. RNA sequencing (RNA-seq) has been successfully used to define the transcriptome and to find novel transcripts absent from the reference genome (Martin and Wang 2011). Compared to genome sequencing, RNA-seq is more economical, especially in the exploration of the complex maize genome containing more than 85% repetitive sequences (Schnable et al. 2009). The construction of the maize pan-transcriptome is especially useful for the discovery of functional dispensable genes. Recently, the maize pan-transcriptome and its diversity have been studied in diverse lines (Hansey et al. 2011; Hirsch et al. 2014), however, we still lack knowledge about many dispensable gene function at the genome-wide level.

Here, with the help of deep RNA-seq of kernels at 15 DAP in a diverse panel with 368 inbred lines (Fu et al. 2013), we characterized the extreme variation at the transcript level (ePAV), relative to the reference genome, and performed association studies between ePAVs and more than 600 quantitative traits. By *de novo* assembly, we also constructed the maize pan-transcriptome and explored its contribution to phenotypic and transcriptomic diversity.

## RESULTS

### Expression presence/absence is prevalent and trans-regulated

Gene expression levels of the annotated genes in the B73 reference genome were quantified using RNA-seq of maize kernels in 368 inbred lines (Fu et al. 2013). We define the expression level differences between subsets of individuals at the given tissue or developmental stage as a polymorphism at the transcription level: expression present/absent variation (ePAV). By filtering the genes showing expression in less than 19 inbred lines or more than 348 inbred lines (MAF≤5%) and applying an adapted distribution-based measure with no subjective set cutoff (see Methods), 13,382 nuclear genes among 38,032 total with ePAVs were obtained (5%≤MAF≤95%) (Table S1). Among them, 6,656 (49.9%) were not explored in a previous study (Fu et al. 2013) since they were expressed in less than 50% but great than 5% of the inbred lines (Fig S1).

Almost half (46%, 6,726) of the ePAV genes expressed in more than 50% of the inbred lines have been clearly identified as regulated by expression quantitative trait loci (eQTLs) in the previous study (Fu et al. 2013). The ePAV genes were more likely to be regulated by distant eQTLs when compared with non-ePAV genes (also called core expression genes, expressed in more than 95% of the lines; *P* < 2.2E-6, χ^2^ test; Fig 1A). The effects of local eQTL were found to be greater than distant eQTL both for ePAV (P= 7.05E-22) and non-ePAV genes (P= 1.92E-135 Fig 1B). The eQTL effects for ePAV genes were greater than those for non-ePAV genes in both local (P=1.34E-18) and distant (P=7.18E-56) types. The ePAV genes were enriched in regulation-related processes (Fig 1C), while the non-ePAV genes tended to play roles as structural genes (Fig 1C). The dominant regulation by distant eQTLs and defined as regulators indicate the ePAVs may act as intermediate regulators to downstream genes, which mostly consist of non-ePAV (or core) genes. This is supported by the observation that most (92%; P<2.2E-16) of the potential regulation targets of ePAV genes were non-ePAV genes. Additionally, the non-ePAV genes tend to be regulated by distant eQTLs as well, with up to 81.2% of regulated non-ePAV genes located on different chromosomes than their ePAV regulators, and for those located on the same chromosome, 86% were separated by over 20Mb (Fig 1D). All the above suggest that the dispensable expression genes are functionally essential, and play key roles in the intermediate regulation layer.

**Fig 1.**
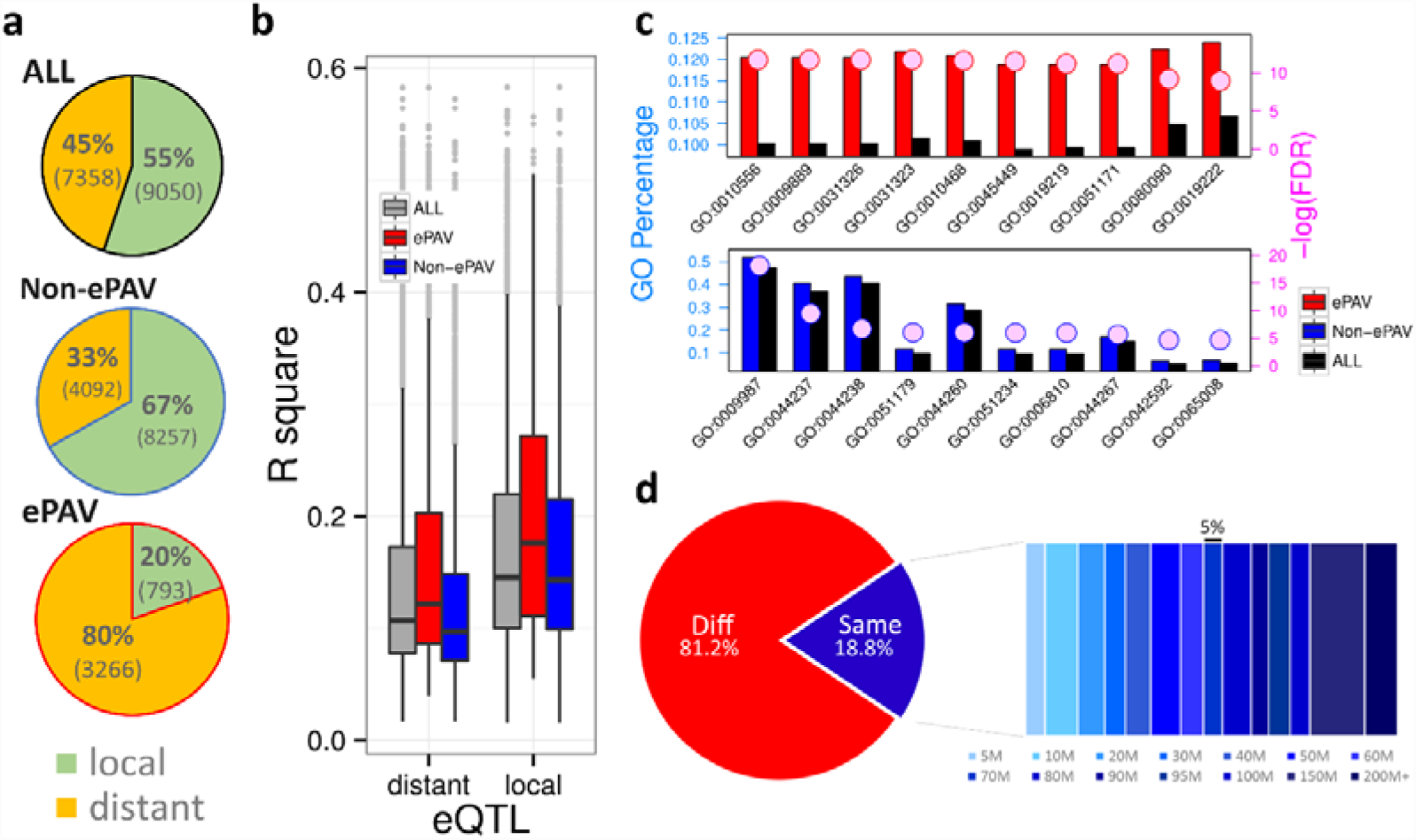
ePAV candidates played key roles in distant-regulation. **(a)** The ratio of local- (green) and distant- (orange) eQTLs among ePAV, non-ePAV and ePAV+non-ePAV together, expressed as percentages. **(b)** The effects of local eQTL were larger than those of distant eQTL both for ePAV (P= 7.05E-22; student test) and non-ePAV genes (P= 1.92E-135). The eQTL effects for ePAV genes were greater than those for non-ePAV genes in both local (P=1.34E-18) and distant (P=7.18E-56) types. **(c)** Top 10 GO enrichment terms in biological processes of ePAV (red) and non-ePAV (blue) are displayed. The left y-axis represents the percentage of genes belonging to each GO term. The colored circles and right y-axis represent the significance level (FDR). Red, blue and black colors means ePAV, non-ePAV and reference levels, respectively. The corresponding GO term description for each GO number: 0010556: regulation of macromolecule biosynthetic process; 0009889: regulation of biosynthetic process; 0031326: regulation of cellular biosynthetic process; 0031323: regulation of cellular metabolic process; 0010468: regulation of gene expression; 0045449: regulation of transcription; 0019219: regulation of nucleobase, nucleoside, nucleotide, and nucleic acid metabolic process; 0051171: regulation of nitrogen compound metabolic process; 0080090: regulation of primary metabolic process; 0019222: regulation of metabolic process; 0009987: cellular process; 0044237: cellular metabolic process; 0044238: primary metabolic process; 0051179: localization; 0044260: cellular macromolecule metabolic process; 0051234: establishment of localization; 0006810: transport; 0044267: cellular protein metabolic process; 0042592: homeostatic process; 0065008: regulation of biological quality. **(d)** ePAV candidates as distant-eQTL affecting expression of Non-ePAV genes. “ Diff” means the eQTL is located on a different chromosome from the gene it regulates and “ Same” both are located on the same chromosome (expressed as %).

### PAV contribute to the causation of ePAV

Before using ePAV for further analysis, we confirmed that the undetectable gene expression in a given tissue was not due to sequencing bias or low sequencing coverage. PAV gene expression should always show ePAV patterns that provide excellent samples to test the reliability of ePAV detection. A ~2.4Mb fragment on chromosome 6 is present in B73 but absent in the Mo17 genome where 62 genes were annotated (Springer et al. 2009). This region was also confirmed by PCR in our inbred lines, among which 209 lines had the same haplotype of B73 and another 15 lines were consistent with Mo17 (Table S2). Among the 62 genes within this region, 61 were detected by RNA-seq and 52 of them were considered as ePAV genes based on the standard (expressed more than 5% and less than 95% lines). The consistency between ePAV status of the 52 ePAV candidates and PCR validation at the genomic level was 74%. This suggests that the ePAV label is acceptable in that most (96.4%) of the inconsistencies were likely caused by non-expression in kernel tissue with presence in DNA sequence, and that the frequency of apparent expression without sequence evidence was rare, at 3.6%.

To determine how many of the ePAVs are caused by genomic PAVs, the reference genome B73 and deep sequenced genome Mo17 were compared. Only 54 (~1%) of the identified 5,838 ePAV genes were supported as sequence PAVs by the re-sequencing results of Mo17 (Fig S2; Lai et al. 2010). However, these two inbreds represent only a fraction of the total maize sequence diversity. Therefore, we used genotyping data generated from Illumina MaizeSNP50 array (50K) for the whole panel (Li et al. 2012) and from the Affymetrix® Axiom® Maize Genotyping Array (600K; Unterseer et al. 2014) for 38 lines; again, we found ~ 1% (102 and 122, or 0.76% and 0.91% for 50K and 600K datasets, respectively) of ePAVs were predicted as PAVs (see Methods). These results together imply that only a small proportion (~1%) of ePAVs were due to PAV in the genomic sequence and therefore, most were likely to be the result of suppression at the expression level.

We further chose 10 putative PAV genes in ePAVs for experimental validation in a subset of 96 inbred lines by genomic PCR. All ten ePAV genes represent genomic PAV genes. The consistency of the ePAV and PAV labels detected by PCR in the 96 lines ranged from 70% to 89% (Fig S3; Table S3), which provided an estimate of the reliability of the predicted ePAV correspondence to sequenced PAVs.

### Novel expressed sequence discovery from *de novo* assembly

RNA-seq reads from each inbred were *de novo* assembled to detect the novel expressed sequences and construct the maize-transcriptome (Table S4). We applied Trinity (Grabherr et al. 2013) to detect novel sequences by comparing the two strategies: “ align-then-assemble” and “ assemble-then-align” (Martin and Wang 2011, detail in methods). Based on the ‘align-then-assemble’ strategy, 7,775 contigs with a total length of 3.46Mb were obtained, of which N50 size was 445bp, much shorter than the average length of reference transcripts (1826bp). Most of these contigs had no hits to protein databases, so it seems that they do not correctly represent the transcripts. We suspect that there was a large proportion of incomplete fragments due to filtering conserved reads and breaking long contigs into short ones. Based on the ‘assemble-then-align’ strategy, 2,355 novel sequences with a total length of 1.9 Mb (N50= 922bp) were obtained (FigS4; Table S5; see methods), resulting in longer and more complete contigs compared to the results of ‘align-then-assemble’ (Fig S5). Further, comparison of the results of the two strategies indicated that some of the sequence reads of conserved functional domains might be filtered out when applying ‘align-then-assemble’ strategy. For example, Unigene_441 from the ‘assemble-then-align’ strategy was identical to Unigene_ref71 from the ‘align-then-assemble’ strategy but longer and containing the unknown protein domain of DUF789. The distribution of unique reads and further PCR re-sequencing both confirmed that the result from ‘assemble-then-align’ was correct (Fig S6; Table S3; Fig S9). Thus, only the results from ‘assemble-then-align’ strategy were used for further analyses.

To evaluate the reliability of the assembled novel sequences, we first compared 2,355 novel sequences to the 4,712 novel genomic contigs obtained in a study of deep sequencing six elite maize inbred lines (Lai et al. 2010), showing that 447 (19%) of our novel sequences align to those novel contigs (Table S6). Second, the novel sequences were compared with 8,681 novel representative transcripts from whole seedling RNA-seq on a panel of 503 diverse maize inbred lines (Hirsch et al. 2014). Nearly 60% (1,380 among 2,355) of the novel sequences identified in the present study had above 85% identity in the alignment with novel transcripts detected from seedling tissue (Table S6). In total, about 62% of our novel sequences were found to have hits in at least one of the previous studies.

We validated the present/absent variation of 10 randomly selected novel sequences in a set of 96 inbred lines including B73 and Mo17 using genomic PCR (Fig S7; Table S3). Two of these novel sequences were present in all 96 inbred lines (Unigene_31, Unigene_361), possibly due to the presence in the genome but absence at the expression level. The other eight were determined to be PAVs, and the consistency of present/absent status between the transcriptome assembly and PAV detected by genomic PCR ranged from 31% to 99%, with an average of 72% (Fig S7). Most (89.2%) of the inconsistency was also due to the presence at the genomic level but with no expression in kernel (Fig S7). We further re-sequenced the amplified products from genomic DNA of the 10 randomly selected novel genes in 5 diverse genotypes and all were consistent with assembly sequences (Fig S8; Fig S9). Cross-comparison with other studies and experimental results not only validates the assembled novel sequences, but also indicates that the predicted present/absent variants are reliable.

### Annotation and mapping of novel expressed sequences

To annotate novel sequences identified in this study, we first compared the sequences with the non-redundant (nr) protein database (Pruitt et al. 2007) using NCBI BLAST, which showed that 1,359 of them had significant matches (E-value < 1e-6) and most (93.57%) of the best matches were within Poaceae. The majority of the significant hits (1,318 of 1,359, 97%) were functionally classified into six types of known enzymes (Fig S10) and conserved domains or annotated motifs (Fig S11; Table S6). In the GO enrichment analysis, the overrepresented processes included several metabolic processes and biotic stimuli (Fig S12; Table S7). In addition, 145 of the 1,037 unannotated novel sequences were considered to have coding potential, having at least 120 amino acid-long predicted open reading frames (ORFs) and a homolog in the non-redundant protein database at a lax standard (*E*-value < 1e-3). Furthermore, 248 of remaining novel sequences were annotated as smRNA precursors against small RNA database (Wang X et al. 2009; *E*-value < 1e-10), and the remaining 644 were predicted to be high confidence novel lncRNAs in maize (Fig S11; Table S6).

To locate possible physical chromosomal positions of the novel sequences, the linkage disequilibrium (LD) mapping strategy was used between novel SNPs within new sequences and high density SNPs in the whole inbred line collection (Fig S13). After multiple sequences alignment, 27,466 SNPs from 664 novel sequences were provisionally identified (See Methods). Based on the LD between SNPs located in novel sequences and high density SNPs with known positions in the whole panel, 625 novel sequences (94.4%) were mapped onto the reference genome (Fig S14; Table S8). The locations of the common expressed genes and the SNPs show the similar trends with enrichment at the ends of the chromosomes, while the distribution pattern of the novel sequences demonstrates fluctuation, and in some cases concentrates near the centromeres on some chromosomes, thus physically complementing the reference genome-containing variations (Fig S14).

### Maize pan-transcriptome plays an important role in regulating phenotypic variation

To systematically explore the genetic consequences of the above described expression variation, and considering that the metabolic phenotype provides a link between gene sequence and visible phenotype, genome wide association study (GWAS) was performed to study the potential effects of ePAV genes on 616 metabolites detected in mature kernels (Wen et al. 2014) and 17 agronomic traits (Yang et al. 2014) measured in the same panel. Among the ePAV genes, 56 (0.42%) were significantly (P < 7.49E-5, 1/n) associated with 15 agronomic traits and 1,967 (14.74%) associated with 526 metabolic traits including content of 18 amino acids (Table S9, Liu et al. 2015).

A major secondary metabolite group in plants, the flavonoids, is widely distributed and has variety of functions (Koes et al. 2005). The pericarp color1 (*p1*) gene encoding an R2R3 Myb-like transcription factor (Grotewold et al. 1994) regulates flavonoid biosynthesis by promoting a suite of structural genes, and conditions pigment in several floral organs (Coe Jr et al. 1988) including the seed coat, cob glumes, tassel glumes, and silk under both genetic and epigenetic regulation mechanisms (Grotewold et al. 1994; Koes et al. 2005; Sekhon et al. 2007; Robbins et al. 2013). In this study, the quantification of 39 flavonoid metabolites together with cob color were used for GWAS, and results indicated the ePAV pattern of the *p1* gene was highly associated with cob color (*P*=1.33E-19), and was correlated with six different flavonoid metabolites (*P*<2.34E-05; Fig 2a). We also identified a structural gene (GRMZM2G162755; anthocyanidin 3-O-glucosyltransferase) significantly associated with cob color (*P*=7.05E-20) and the same six flavonoid metabolites (*P*<6.23E-05; Fig 2a). This gene was shown to be regulated by *p1* in a previous eQTL mapping study (Fu et al. 2013) and ChIP-Seq analysis (Morohashi et al. 2012). Another copy of the R2R3 Myb-like transcription factor (*p2*, GRMZM2G057027) could regulate the other two flavonoid metabolites (*P*<3.41E-06; Fig 2a). Notably, we found these three ePAV candidates, as regulators, could also control expression of other genes that are related to flavonoids such as *c2*, *chi1*, *a1*, *pr1* and *whp1* (Grotewold et al. 1994; Morohashi et al. 2012; Fig 2b; *P*≤9.65E-10), providing further support for the hypothesis that these ePAV genes are involved in the flavonoid pathway and functioning through the PAV differences in expression level.

**Fig 2.**
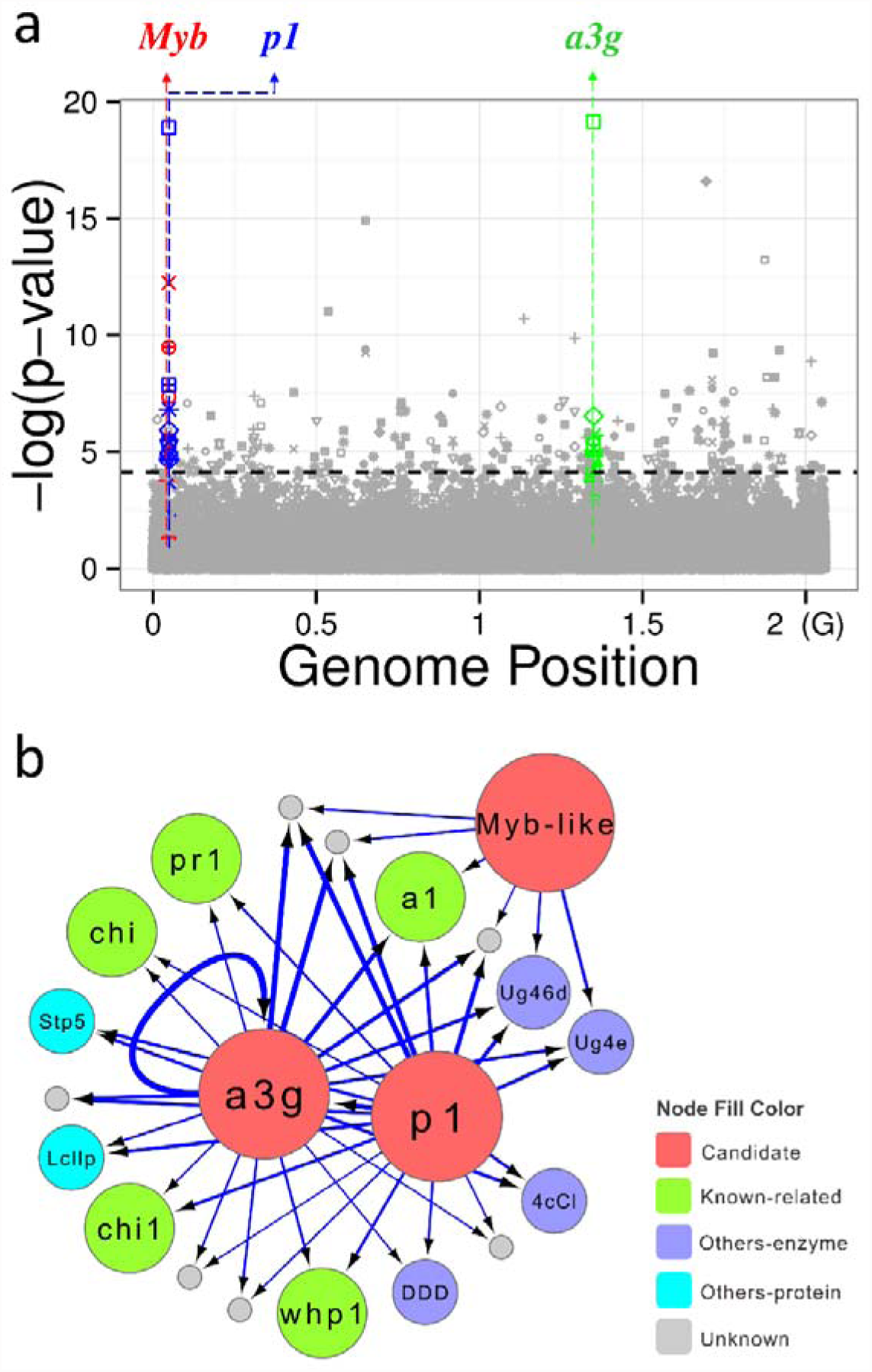
ePAV candidates contributed to both maize cob color and various kinds of flavonoids. **(a)** Manhattan plot of the association of three ePAV candidates, maize cob color and several flavonoids. Different shapes represent different traits, and points with different color represent different kinds of ePAV candidates: Blue: *pericarp color1* (*p1*, GRMZM2G084799); Red: *p2*, another copy of R2R3 Myb-like transcription factor (GRMZM2G057027); Green: anthocyanidin 3-O-glucosyltransferase (GRMZM2G162755); Grey: other ePAVs. Black dashed horizontal line was the cut-off (P=7.47E-5) of significant level. **(b)** The three ePAV candidates were also significantly associated with expression of related genes within maize flavonoid pathway. Nodes in red are the three ePAV candidates above, green nodes represent several identified genes located in the maize flavonoid pathway, purple nodes are other genes encode enzymes, light blue were other genes encoding non-enzyme proteins (such as transporters), and grey nodes had no annotation. The blue arrow edges link the ePAV candidates and its associated targets and the a3g links to itself meaning self-regulation in expression level, while thicker lines represented more significant associations.

The novel expressed sequences play critical roles in regulation of the transcriptome and metabolome. For the novel sequences, a re-mapping strategy was applied to correct the PAV distribution for each novel gene, to be used in GWAS (see Methods). We found that 26 (1.1%) of the novel genes were associated (P < 4.25E-4) with 13 agronomic traits (Table S9). Eleven were associated with flowering time (i.e. Days to Tasseling, Days to Pollen Shed and Days to Silking). We also identified a novel gene (Unigene_55) that encodes a late embryogenesis abundant (LEA) protein that is associated with kernel width (P=2.14E-5). LEA proteins have been described as accumulating late in embryogenesis and could protect other proteins from aggregation under various environmental stresses (Goyal et al. 2005). Here we provided a clue that LEA may also affect kernel size.

Moreover, 788 novel genes (33.46%) were associated (P < 4.25E-4, 1/n) with 487 metabolic traits measured in maize kernels (Table S9, Wen et al. 2014), which implied that those novel genes could play more complex roles within cellular metabolism processes; thus, this study provides fresh resources for the genetic study of maize kernel quality and production. Metabolic processes are commonly controlled by transcription regulation. Therefore, it is valuable to examine whether the identified novel genes were widely involved at multiple regulatory levels. We found the novel sequences were significantly responsible for expression levels of annotated genes or their expression presence/absence states at a strict cutoff (*P*≤1E-4), including the novels annotated as non-coding RNAs (Table S9). By combing the metabolome and transcriptome findings, 23 novel genes were found playing roles in both metabolic processes and expression regulation.

Plant NBS-LRR proteins can directly or indirectly recognize pathogen-deployed proteins and triggers plant defense responses (McHale et al. 2006; DeYoung and Innes 2006), and exhibit high levels of PAV polymorphism in various plants (Shen et al. 2006; Yang et al. 2008; Tan et al. 2012; González et al. 2013; Wu et al. 2014). Here, we identified two novel NBS-LRR genes (Fig 3a, b) that have high homology to rice NBS-LRR genes Os11gRGA4 and Os11gRGA5, which were shown to interact functionally and physically to mediate resistance to the fungal pathogen *Magnaporthe oryzae* (Okuyama et al. 2011;Césari et al. 2014). Recent studies have verified that inducing plant immunity impacts flavonoid biosynthesis (Ali et al. 2011; Serrano et al. 2012), and that flavonoid compounds significantly contribute to plant resistance (Treutter 2005; Treutter 2006).Os11gRGA4 was found to be associated with the flavonoid Naringenin O-malonylhexoside (Chen et al. 2014). Interestingly, we found that these two novel NBS-LRR like sequences were both associated with Apigenin C-pentosyl-O-coumaroylhexoside and C-pentosyl-apigenin O-caffeoylhexoside contents, two flavonoid metabolites (Fig 3c). In addition, these two novel sequences were also associated with several gene expression presence/absence states (Fig 3d), including transcription factors with DNA binding activity (TGA6 or GRMZM2G000842; GRMZM2G405170), spliceosomal complex (GRMZM2G011034), nucleic acid binding genes (GRMZM2G088348), translation release factor (GRMZM5G864412), actin cytoskeleton (GRMZM2G552644) and other enzymes functioning in metabolic processes. These observed associated targets were consistent with previous observations that alternative splicing is important in the regulation of NBS-LRR proteins and plant immunity (Jordan et al. 2002; Zhang et al.2014), and that TGA6 and other bZIP transcription factors are significant in plant defense against pathogens (Xiang et al. 1997; Alves et al. 2013). Actin cytoskeleton dynamics also play an important role in mediating resistance (Wang et al. 2013), and translation release factors are critically involved in the elimination of aberrant mRNA. The regulatory targets in pathogen defense response are indeed R-genes, especially for the abundant and alternatively spliced NBS-LRR R-genes (Riehs-Kearnan et al. 2012).

**Fig 3.**
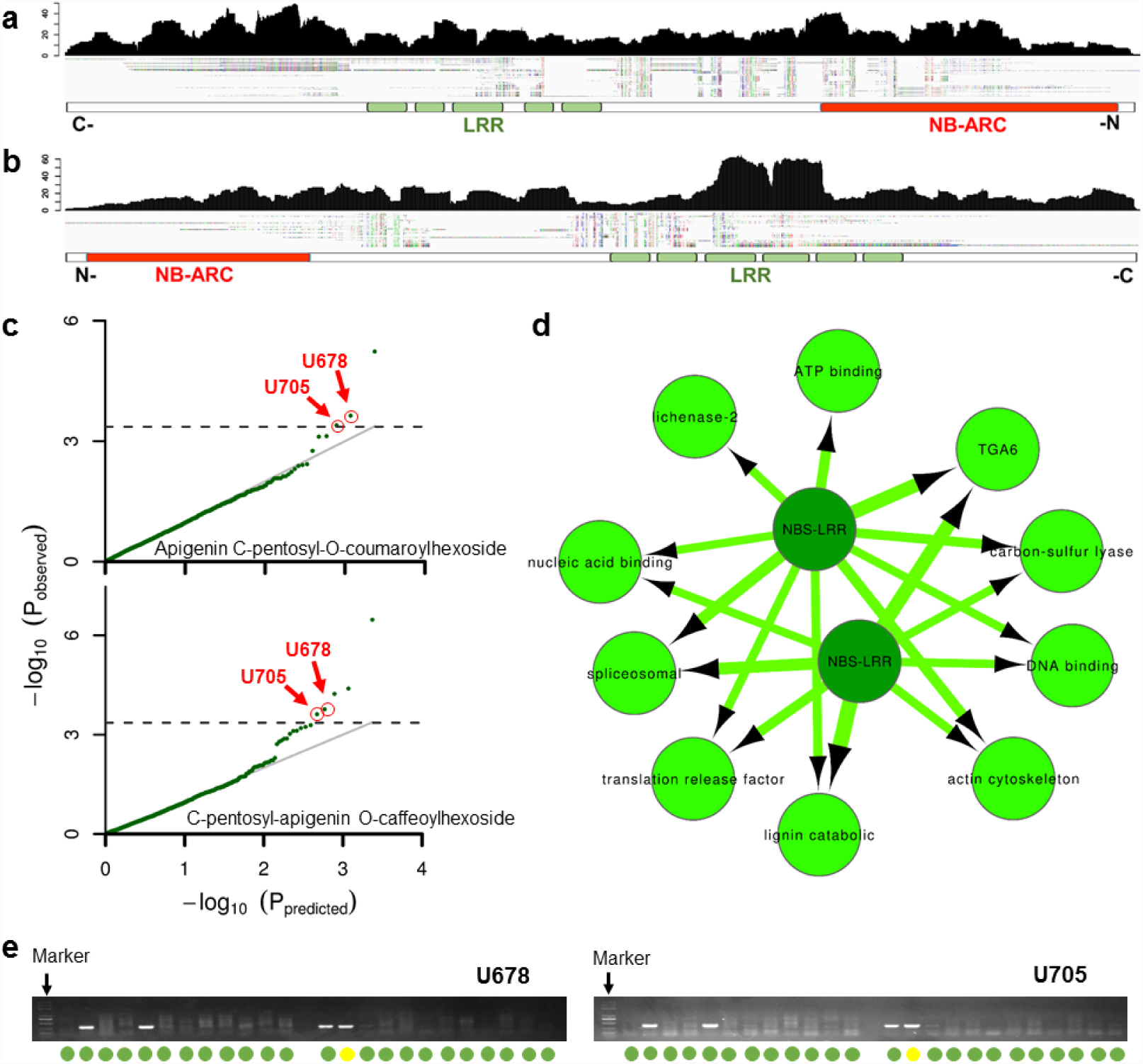
Two novel NBS-LRR genes showed significant association with flavonoid metabolites and with expressed genes involved in flavonoid pathway. Read distribution and predicted conserved domains of novel reference gene Unigene_678 (a) and Unigene_705 (b) and sequence alignments for all presence genotypes. (c) Q-Q plot of association mapping for different flavonoids. (d) The two novel NBS-LRR genes were also significantly associated with other genes with expression presence-absence variation. (e) Validation of the PAV of the two novel genes. Green represents consistency between experiment and prediction. Yellow means the gene was absent in our prediction but exists in the genome.

The two novel sequences were confirmed by PCR sequencing (FigS9; Table S3). Moreover, the consistency of presence/absence variation in the association panel used in our study between observed and predicted variants was greater than 98% (Unigene_678) and 96% (Unigene_705), respectively. The PAV states of these two genes on the genome level also showed a significant relationship with the metabolic traits mentioned above. These results show that the dispensable novel expressed sequences were important both in morphological adaptation processes as previously reported (Hirsch et al. 2014), and in cellular metabolome and transcriptome regulation.

### Present and absent genes may contribute to trait heterosis

Complementation of gene content variation is assumed to be important in heterosis (Fu and Dooner 2002; Springer et al. 2009; Lai et al. 2010; Schnable and Springer 2013). Since our identified expressed novel genes have been shown to be functionally important, their combination of inbred-specific sequences in hybrids could provide novel trans-interactions potentially resulting in non-additive expression. This provides an opportunity to test the link between gene content PAV and heterosis. We crossed the association panel with the Mo17 inbred line to develop a suitable population to test this. Six yield-related traits were measured for each hybrid in different environments over two years (Methods). The degree of heterosis increased with more complementary (present in one and absent in the other inbred parent) novel genes in the hybrids among five of the six measured traits (Fig 4A). This trend is more significant for those traits with relatively stronger heterotic effect, and novel sequences identified in this study have a greater effect than ePAVs (Fig 4B). However, only a small portion (<10%) of observed heterosis was explained by novel sequences and/or ePAVs, which implies that heterosis is complexly affected by many different factors (Stupar and Springer 2006; Swanson-Wagner et al. 2006; Lai et al. 2010; Hansey et al. 2012; Guo and Rafalski, 2013).

**Fig 4.**
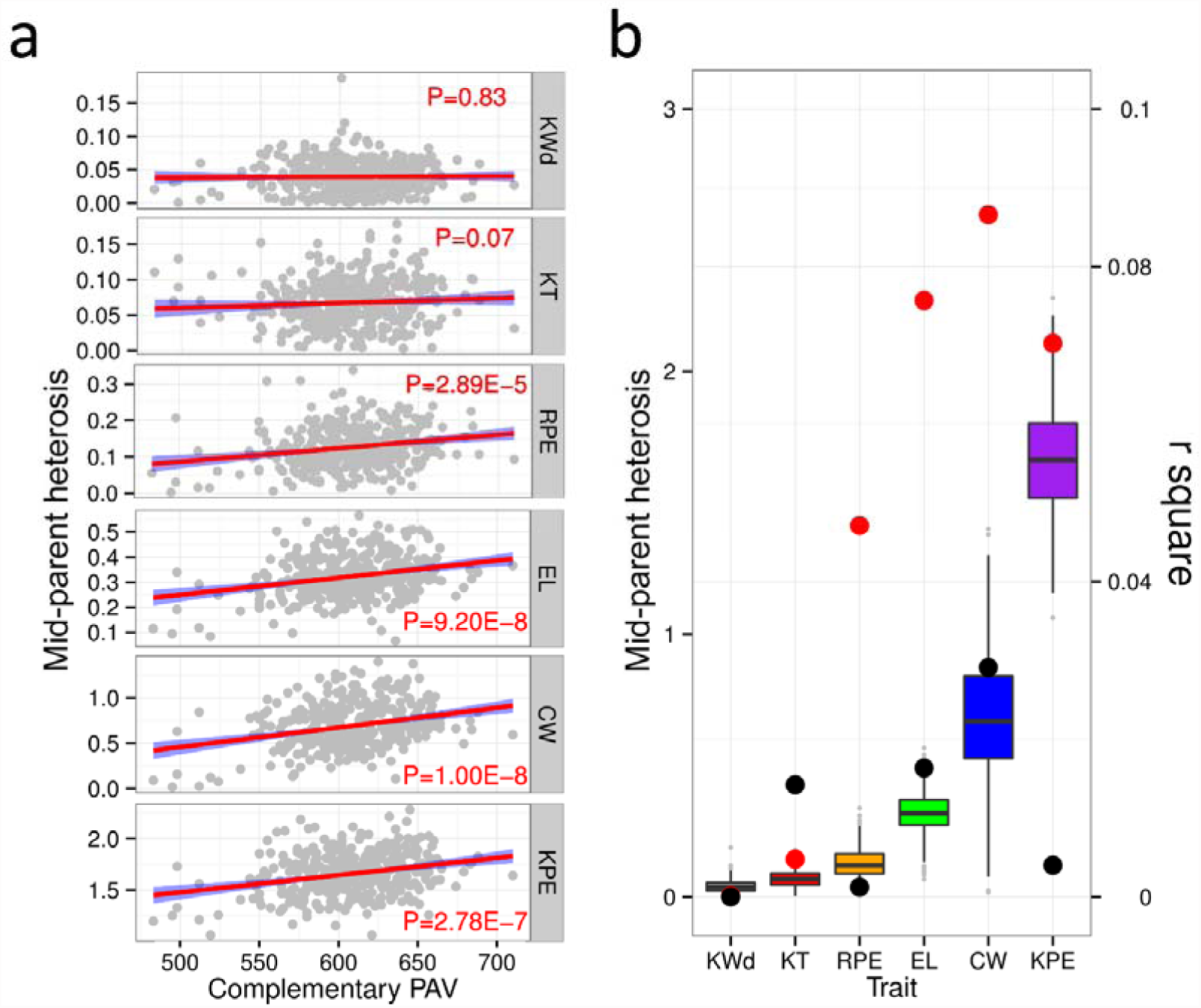
PAV status of novel genes and ePAV both correlated with heterosis of most yield-related traits. **(a)** Correlation between mid-parent heterosis and the number of complementary novel genes exhibiting PAV between the parents of the F1 population. Six panels represent different yield-related traits. KWd: Kernel Width; KT: Kernel Thick; RPE: Rows Per Ear; EL: Ear length; CW: Cob Weight; KPE Kernels Per each row of Ear. **(b)** Boxplot in different colors represents different traits ordered by mid-parent heterosis (the left y axis). The points in red and black represent Pearson’ s r^2^ of correlation between mid-parent heterosis and the number of complementary novel genes and ePAVs showing presence-absence variation between parents of the F_1_ population.

## DISCUSSION

### The importance of ePAV

Expression PAV is a kind of variation at transcript level, mostly due to genetic or epigenetic regulation. With the conservative distribution-based approach (see Methods), we identified more than 13,000 genes as ePAVs, about one third of the maize annotated genes. These are genes that are only expressed in a subset of the association panel. This finding was based on one tissue (kernel of 15DAP) and limited inbred lines (n=368), therefore, more ePAVs will be identified when more tissues and materials are studied. The number of ePAV should vary under different sequencing coverage and even different cutoffs; however, this study demonstrates that ePAV is a common phenomenon and that the underlying mechanisms and ramifications need to be explored. Among our findings, the ePAV genes were enriched in regulation-related processes and were usually regulated by distant eQTLs while core expression genes were commonly regulated by local eQTLs (Fig1 a). The different regulatory patterns imply that the two kinds of genes may affect phenotypic variation by different mechanisms.

Transcript variation as an independent variable can be regarded as a molecular marker to perform GWAS (i.e. ePAV-GWAS), corrected for population structure and relatedness, and this should provide additional insights into the architecture and regulation of quantitative traits and help understand certain important biological questions such as adaptation (Hirsch et al. 2014; Harper et al. 2012). Interestingly, about 15% of the identified ePAVs were found to associate with agronomic and metabolic traits, which confirms the expectation that gene expression presence and absence can affect the phenotypic variation directly. Combining the ePAV-GWAS results with SNP-GWAS and eQTL mapping information aid not only in the identification of gene candidates, but also to better understand molecular mechanisms. Here, we use cob color and several related flavonoid metabolites as an example for further exploration. After strict filtering of our genotypic data, there were no SNPs left within the *p1* locus (Fig S15b), known to control cob color, but an associated region was found upstream of the the *p1* locus (~200K, Fig S15a,b). This makes it difficult to unambiguously identify a single causal gene for cob color. Using ePAVs as markers (ePAV-GWAS), the *p1* locus was exactly identified and the *p2* was promoted as another candidate (Fig2a, Fig S15b, panel4). Another significant locus (GRMZM2G162775, a3g) on chromosome 6 was detected by ePAV-GWAS (Fig S15a, panel4), and may have been detected as regulated by *p1* by the previous eQTL mapping (Fig S15a, panel c; Fu et al. 2013) and ChIP-Seq studies (Morohashi et al. 2012). This trans-regulation pattern could not be discovered by applying SNP-GWAS, even when the SNP density was doubled to 1.25 million (data not shown). Thus, the ePAV will provide a unique complementarity to SNP marker for the genetic exploitation.

### The “dispensable” genes are important functionally

Although more than 13,000 ePAVs were identified, only small proportion (~1%) included genomic PAVs which are sometimes called dispensable genes. Previous studies revealed that the B73 reference genome included only 70% of the total low-copy sequences available in the maize species (Gore et al. 2009), which implied that many dispensable genes are present beyond the reference genome. It is necessary to explore these novel genes that may be phenotypically important in certain genotypes. We applied *de novo* assembly to detect such novel sequences. Of the two combined strategies were available ‘assemble-then-align’ resulted in longer and more complete contigs. Although in theory, ‘align-then-assemble’ should be more sensitive and *de novo* assembly was likely to work only for the most abundant transcripts (Haas and Zody 2010), in practice, the align-first strategy probably enrich some low abundance (and possibly extraneous) reads and assemble them into contigs. Only a small portion (4%, identity≥85%, coverage≥85%) of the contigs from the align-first strategy could identify high confidence matches in assemble-first contigs. After stringent filtering, 2,355 high confidence novel sequences with a total length of 1.9Mb were obtained.

The enrichment analysis of these novel sequences suggested their roles in metabolic processes responsive to stimuli. Almost 34% of them were found to be associated with metabolic traits. The differences in metabolism - related genes may be associated with differential environmental effects (Liu et al. 2015). The novel sequences involved in development, such as beta-tubulin, also likely contribute to adaptation. In hybrids, the number of novel genes in heterozygous (present in one parent, absent in the other) state was correlated with heterosis of yield-related traits, which supports the complementation hypothesis of this phenomenon. We showed that the “dispensable” genes, whether they were present on reference genome or not, indeed play an indispensable roles at the population level.

### The size of maize pan-transcriptome

The construction of the maize pan-transcriptome is more effective than a maize pan-genome because the high proportion of repetitive sequences present in the genome complicates assembly. However, limitations in tissue and availability of diverse genotypes could result in underestimating the size of maize pan-transcriptome (Fig S16a). We compared tissue-specific and genotype-specific efficiency in discovering novel transcripts. We selected five diverse tissues (16DAP Whole Seed, V3_Stem and SAM, V9_Immature Leaves, R2_Thirteenth Leaf, 6DAS_GH_Primary Root) from a previous study (Sekhon et al. 2013) and repeated the *de novo* assembly process as described. We found more novel transcripts when adding new tissues than by adding new individuals (Fig S16b). This indicated that the expression divergence is significantly larger between tissues than individuals (P=1.07E-58).

We estimated the size of maize pan-transcriptome based on our RNA-seq data from one tissue but multiple genotypes (Fig S16c, see Methods). As expected, when adding more genotypes to the analysis the number of additional novel sequences detected eventually leveled off and the total is expected to reach an asymptotic maximum of ca. 28,000. Using the reference genome and a similar procedure, we found that number of core expression genes decreased and became nearly invariable when more than 200 lines were included. The minimum number of core expressed genes and maximum for dispensable ones were 22,043 and 13,382, respectively (Fig S16d). Combining the reference based genes and newly identified genes from the current study, we estimated the size of the pan-transcriptome of the maize whole kernel is about 63,000. Under the simple assumption that maize kernels only express 70% ~ 80% of the total genes (Fu et al. 2013), the whole pan-genome of maize is close to 78,000 ~ 84,000. Thus, the present reference genome may only capture half of the predicated maize pan-gene, which is similar to previous prediction (Hirsch et al. 2014). To identify the pan-gene and study the functions will help to understand the genome better thus enhancing crop improvement.

## METHODS

### Detection of ePAV

To quantify the expression of 38,032 reference genes, read counts for each expressed gene and individual transcripts of that gene were calculated and scaled according to the definition of RPKM (reads per kilobase of exon model per million mapped reads). The genes showing expression (RPKM=0) in less than 19 (5%) inbred lines were excluded in the following analysis. In addition, we filtered the genes which expressed (RPKM>0) in more than 348 (95%) inbred lines. The remaining genes were considered to have presence/absence variation in expression, and several further distribution-based steps were used to acquire an ePAV pattern: (1) extract non-zero expression data of a ePAV gene in 368 inbred lines; (2) sort it from smallest to largest and make the frequency distribution (10 groups); (3) turn the abnormal low (data in 1st group) and high (data in groups that frequency<3) expression values according to the frequency distribution as “ NA” (4) convert the rest of no-zero expression data to ‘1’ and no expression data to ‘0’ to get the ePAV pattern of each gene.

### Prediction of PAV through genotyping from 50K and 600K SNP arrays

The ePAV genes (including 1Kb upstream and downstream regions) containing at least two SNPs in the array genotyping dataset were used to analyze their PAVs. A gene was regarded as potential PAV if all its SNPs were genotyped as missing in a particular line. Further, for each ePAV, if the potential PAV occurred at more than a certain ratio (5%, or 19 for 50K and 2 for 600K datasets, respectively), it was considered as non-random, thus to be candidate PAV.

### *De novo* Transcript Assembly

The poly(A)^+^ transcriptomes of immature kernels (15 DAP) were sequenced using 90-bp paired-end Illumina sequencing with libraries of 200-bp insert sizes. The sequencing data for this project can be downloaded in the NCBI Sequence Read Archive under accession code SRP026161. Average 73.9 million reads were obtained in each sample and 367 inbred lines were used in assembly process (Table S4; Li et al. 2012; Fu et al. 2013). The adaptors and low quality reads were filtered using Trimmomatic software (Bolger et al. 2014), resulting in a total 24.7 billion high-quality reads, used for the assembly (Table S4; Fig S17).

While applying the “ align-then-assemble” strategy, the mapping process was first performed by Bowtie2 version 2.0.2 (Langmead et al. 2012) and TopHat2 version 2.0.6(Kim et al. 2013) with the parameters -i 5, -I 60000, -r 20, --mate-std-dev 75 and gene annotation was provided. The unmapped reads from each individual were assembled by Trinity (Grabherr et al. 2011), which is based on the de Bruijn graphs algorithm. Min count for K-mers to be assembled by Inchworm most influenced the result. We found that there was a large increase in the numbers of transcripts that align to the B73 reference transcripts when applying min K-mers between 2 and 3 and we chose the parameters: -seqType fq,-min_kmer_cov 2, -min_contig_length 200. In the “ assemble-then-align” strategy, the whole cleaned RNA-seq reads from 367 inbred lines were *de novo* assembled with the same parameters.

### Identification of Novel Sequences

To detect truly novel sequences and remove the ones that were alleles or paralogs of sequences present in the B73 reference genome, the assembled transcripts from each line were aligned to B73 5b pseudomolecules using GMAP (Wu and Watanabe 2005), a genome alignment program for mRNA sequences. We randomly chose 200 assembled transcripts from each inbred line to determine GMAP parameters and the identity cutoffs were then set to 0.85. The representative transcripts that did not align to the reference sequence were clustered by the TGI Clustering tool (TGICL; Pertea et al. 2003). Transcripts present in at least 19 inbred lines (5% of 367) with non-homology (identity < 95%; coverage < 90%) to B73 cDNA 5b pseudomolecules (FGS) were retained as candidate novel sequences. DeconSeq (Schmieder and Edwards 2011) was further used to remove sequence contamination to improve the reliability of novel sequence identification. Reference genomes of human, human microbiome, and virus were used as the “contamination-datasets”, and plant datasets including *Zea mays, Oryza sativa, Sorghum bicolor, Setaria italica*, and *Brachypodium distachyon* were used as “ retain-datasets” under the parameters of identity≥98% and coverage≥90%. Finally, RNA-seq reads were aligned back to novel sequences for quality assessment by running alignReads.pl in Trinity software (Grabherr et al. 2011) with the --bowtie and --phred64-quals options. The 12 sequences which had breakpoints in distribution of reads were excluded. This indicates there may be minor errors in the transcripts assembly process. All the procedures and related results were shown in Fig S17. On average, 57,628 assembled transcripts with N50 size 1,078 bp were obtained for each inbred line (Table S4). After excluding the transcripts present in the B73 reference and other contaminations, an average of 1,388 unmapped transcripts were retained in each inbred line. We clustered these remaining transcripts from all inbred lines, and the longest one was selected as a representative sequence in each unigene cluster. Each unigene cluster would then be retained if it was present in at least 19 inbred lines (5% of the panel). Finally, 2,355 novel representative sequences with a total length of 1.9 Mb and N50 size 922bp were obtained (Fig S4; Table S4).

### An improved re-mapping strategy to correct the distribution of novel sequences

After the clustering step, we obtained the PAV patterns for each novel gene among all genotypes. When considering those present in genomic sequence but non-expressed as “ inconsistent”, the consistent ratio reached to average 67%, using a simple clustering step. However, we found that some novel genes appear to be also expressed in predicted “ Absence” lines

This may be caused by incorrect assignment to the genomic location of short or well-conserved expressed sequences. In these cases we applied a second re-mapping step to recover correct genomic matches, by using BLASTx with “ identity≥0.96, query-coverage≥0.5, subject-coverage≥0.96”, which improved the consistency ratio to an average of 72%.

### Annotation of Novel Sequences

Blast2go (Conesa and Götz 2008) is an all in one tool for functional annotation of novel sequences and the analysis of annotation data. For BLASTx to nr database (Pruitt et al. 2012), a minimum E-value of 1e-6 was used and only best hits were considered. BLAST XML result file was imported in Blast2go. GO mapping and InterProScan (McDowall and Hunter 2011) were also performed to complete the annotation. A total of 1,359 novel sequences had matches in the nr protein database using BLASTx (*E*-value ≤1e-6). Nearly all of them (1,318 of 1,359, 97%) can be functionally classified into families and contained conserved domains and functional sites (Table S6). The remaining 1,037 unannotated sequences were left. Among annotated ones, 166 could encode enzymes (Fig S10; Table S6). 640 can be grouped into at least one GO term (Table S6) and used in the next GO enrichment analysis.

The remaining unannotated novel sequences were used to predict the protein coding potential. Three important criteria were used: transcript length, open reading frame (ORF) size, and presence of homology with known proteins. Transcript length was set to 200bp. Only three ORFs longer than 120 amino acid were identified in 14 known long non-coding RNAs (lncRNAs; Boerner and McGinnis 2012), ORFs longer than 120 amino acid considered potential coding candidates. Coding regions and the corresponding amino acid sequences were extracted from novel sequences using TransDecoder in the Trinity software (Grabherr et al. 2011). In addition, transcripts were aligned to UniProtKB/Swiss-Prot database (Jungo et al. 2012) identify transcripts with potential protein-coding ability (E-value≤1e-3). The unaligned transcripts were considered non-coding RNAs (ncRNAs). Using BLASTN (E-value≤1e-10) and Infernal software (“INFERence of RNA ALignment”; Nawrocki and Eddy, 2013; score≥40), 248 sequences were matched to NONCODE database (Bu et al. 2012), smRNA transcriptome databases including predicted microRNAs (miRNAs), other predicted short hairpin forming RNAs (shRNAs) and predicted small interfering RNAs (siRNAs) or Rfam database (Wang X et al. 2009; Burge et al. 2013). These sequences were all considered the precursors of small RNAs. The remained 644 novel sequences were predicted to be high confidence maize lncRNAs (Fig S11; Table S6).

### SNP Analysis and LD Mapping of Novel sequences

MAFFT (Katoh et al. 2009) was used to align novel sequences from all inbreds to their corresponding representative ones (the longest one in each unigene cluster; see above). SNP_SITES software (<u>https://github.com/sanger-pathogens/snp_sites</u>) was used to identify SNPs in the multiple alignment. Biallelic SNPs with minor allele frequencies (MAFs) larger than 0.05 were retained for analysis. A total of 27,466 SNPs were identified in 664 novel sequences. Pairwise LD between SNPs within novel sequences and between SNPs in B73 reference genome was computed by a script on the assumption of equal probability for either phase relationship of the alleles. B73 reference gene with the highest LD (and at least *r*^2^>0.1) to SNPs within the novel gene was considered the likely location of the novel gene. Using this approach, of the 664 novel sequences with SNPs, 627 were mapped to the B73 reference.

### Validation of PAVs within ePAV and novel ones

Genomic PAV for 10 ePAV genes, and the 10 novel expressed genes across a set of 96 diverse inbred lines were evaluated using touchdown PCR. Inbred lines information, primer sequences and experiment results are available in Fig S3,Fig S7, Table S3. The thermo cycler program for touchdown PCR were included: 1=94°C 5min; 2=94°C 30s; 3=64°C 30s -0.5°C/cycle; 4=72°C 50s; 5=GOTO 2 12repeats; 6=94°C 30s;7=58°C 30s; 8=72 °C 50s; 9=GOTO 6 23repeats; 10=72 °C 5min;11=25°C 2min; 12=END. The PCR products of 10 novel genes in 5 lines were then re-sequenced and subjected to multiple alignment to evaluate the correctness of *de novo* assembly.

### Novel sequences and ePAVs both contributed to heterosis

To test whether the PAV pattern of the novel genes contributes maize heterosis, all population inbred lines were planted with randomized complete experimental design by single replication in 2011 (Chongqing city; Hebi city, Henan province; Honghe autonomous prefecture, Yunnan province and Sanya city, Hainan province) and 2012 (Chongqing city; Hebi city, Henan province; Honghe autonomous prefecture, Yunnan province and Wuhan city, Hubei province) and 6 yield-related traits including Kernel Width (KWd), Kernel Thickness (KT), Rows Per Ear (RPE), Ear Length (EL), Cob Weight (CW), Kernels Per Ear (KPE) were measured. Generally, the average values from five individuals were calculated to represent each line in each experiment and the BLUP values from different environments and years were used for next analysis. The number of PAVs in heterozygous state (present in one parent, absent in the other) were then used to evaluate their correlation (R-square was measured; Fig 5) with observed mid-parent heterosis for each trait.

### The estimation of the maize pan-transcriptome size

Five diverse tissues (16DAP Whole Seed, V3_Stem and SAM, V9_Immature Leaves, R2_Thirteenth Leaf, 6DAS_GH_Primary Root) from a previous study (Sekhon et al. 2013) were chosen to repeat *de novo* assembly process. We then compared each pair of tissues and individuals, by measuring the ratio of shared genes to total genes. To eliminate the effect of biased sample size (5 tissues vs 367 individuals), we randomly selected five pairwise comparisons and repeated this process 1000 times, then compared resulting distribution.

To determine whether the maize pan-transcriptome is open (the size of pan genome grows continuously with the number of sequenced individuals increases) or closed (the size of pan genome reached a constant value with the number of sequenced individuals increases) and to estimate the size of it, a simulation process on real data was used. There were three parts to form the maize pan-transcriptome: reference based core genes, reference based dispensable genes (ePAV) and novel sequences. We randomly chose 20 samples in 367 maize inbred lines in a clustering run to estimate the number of novel sequences among them and then add another 20 lines to do the same cluster recursively until a total of 360 inbred lines were in the set. Ten independent simulations were run and the mean of each run from n = 20 to 360 inbred lines was used to estimate the maximum number of novel sequences. The same simulation process was also performed to estimate the maximum value of core genes and dispensable genes on references genome.

## DATA ACCESS

The raw RNA sequencing data have been deposited in NCBI Sequence Read Archive (SRA) under accession SRP026161. The raw sequences (with fasta format) and annotation information of novel assembled ones could be available at www.maizego.org/Resources.

## ACKNOWLEDGMENTS

This research was supported by the National Natural Science Foundation of China (31123009 and 31222041) and the National Hi-Tech Research and Development Program of China (2012AA10A307) and the National Youth Top-notch Talent Support Program.

## AUTHOR CONTRIBUTIONS

J.Y. and J.F. designed and supervised this study. M.J., H.L. and C.H. performed the data analysis. Y.X. provided the simulation code for pan-genome size estimation. G.W. and W.X. contributed to materials collection and suggested analysis procedure. M.J. and Y.W. performed the experiments. M.J., H.L. J.Y. and J.F. prepared the manuscript, and all the authors critically read and approved the manuscript.

